# Deciphering of BTH-induced response of tomato (*Solanum lycopersicum* L.) and its effect on plant virus infection through the multi-omics approach

**DOI:** 10.1101/2022.07.08.499279

**Authors:** Frąckowiak Patryk, Wrzesińska Barbara, Wieczorek Przemysław, Sanchez-Bel Paloma, Kunz Laura, Dittmann Antje, Obrępalska-Stęplowska Aleksandra

## Abstract

One of the preventive methods used to limit the losses caused by viruses is the application of synthetic immunity inducers, such as benzo(1,2,3)-thiadiazole-7-carbothioic acid S-methyl ester (BTH). This study aimed to explain how the BTH treatment affects the defence and developmental processes in tomato plants (*Solanum lycopersicum* L.) as well as plant response to virus infection.

The comparative multi-omics analyses of tomato plants treated with BTH were performed, including transcriptomics (RNA-seq), proteomics (Liquid Chromatography-Mass Spectrometry), and metabolomics (targeted hormonal analysis). To confirm the priming effect of BTH on tomato resistance, the plants were infected with tomato mosaic virus (ToMV) seven days post-BTH treatment.

The combined functional analysis indicated the high impact of BTH on the plant’s developmental processes and activation of the immune response early after the treatment. In the presented experimental model, the increased level of *WRKY TRANSCRIPTION FACTORS, ARGONAUTE 2A*, thiamine and glutathione metabolism, cell wall reorganization, and detoxification processes, as well as accumulation of three phytohormones: abscisic acid, jasmonic-isoleucine (JA-Ile), and indole-3-carboxylic acid (I3CA), were observed upon BTH application.

The immune response activated by BTH was related to increased expression of genes associated with the cellular detoxification process, systemic acquired resistance, and induced systemic resistance as well as post-transcriptional gene silencing. Increased levels of I3CA and JA-Ile might explain the BTH’s effectiveness in the induction of the plant defence against a broad spectrum of pathogens. For the first time, the BTH impact on the thiamine metabolism was revealed in tomatoes.

## 1. Introduction

Tomato (*Solanum lycopersicum* L.) is one of the most widespread vegetable crops, important in the human diet, and a rich source of minerals, antioxidants, carotenoids (mainly β-carotene) and vitamins (Yusufe et al. 2017). Exposed to a wide array of stresses, including abiotic and biotic factors, the tomato plants have developed various mechanisms of response to ubiquitous stimuli (Santner and Estelle 2009; Verma et al. 2016). Plant hormones (phytohormones) are responsible for maintaining metabolic balance in plants, from the processes of growth to those of defence response. Particularly important phytohormones associated with this equilibrium are salicylic acid (SA), jasmonic acid (JA), abscisic acid (ABA), and auxin (Aux/IAA) (Denancé et al. 2013). The whole-plant signalling pathways are interconnected. The sophisticated interaction between hormones and genes related to each signalling pathway enables plants to survive unfavourable conditions.

The metabolic homeostasis of plants can be compromised by various environmental factors, including pathogens, among other viruses (Suzuki et al. 2014). Importantly, besides prevention, there is no other method to control virus spread. To this aim, a combination of chemical and biological methods supported by agricultural technologies is being used (Maksimov et al. 2020). The best approach to minimize the risk of crop losses caused by viruses is, inter alia, the use of the resistant plant varieties in the production, the eradication of diseased plants from the field, and control of the population of virus vectors, among others insects (Nicaise 2014). An important group of prevention agents reducing the negative effects of virus infection are natural or synthetic immunity inducers, such as chitosan (Chirkov 2002) or thiadiazoleacetamides, for instance, benzo(1,2,3)-thiadiazole-7-carbothioic acid S-methyl ester (BTH) (Zhao et al. 2006) or its derivatives. Resistance inducers mobilize the plant immune system, and most importantly, are not phytotoxic (Pospieszny 2017). However, activation of the plant ‘immune system’ by resistance inducers is often associated with a fitness cost (Trejo-Saavedra et al. 2013).

Pathogens trigger a wide range of defence mechanisms in the plant, for instance, changes in the cell wall composition, activation of oxidative burst, synthesis of resistance compounds, or up-regulation of defence-related genes, among others those encoding pathogenesis-related proteins (PRs) (Dodds and Rathjen 2010). The BTH, a functional analogue of SA, activates the same pathways as in the pathogen response and initiates the induction of systemic acquired resistance (SAR) in the treated plants (Smith-Becker et al. 2003). The first use of BTH in modern agronomy dates back to 1996 (Thomson et al. 1998). Recently, research on BTH has been intensified and focused on its mechanisms of action and effectiveness in various biological systems. Noteworthy, new chemically modified BTH derivatives are being synthesized to improve their effectiveness through better assimilation by plants, higher dissolution rate, biodegradability, and protective potential (Smiglak et al. 2016, 2017).

The BTH-mediated resistance induction was evaluated, among others, in pepper against pepper golden mosaic virus (Trejo-Saavedra et al. 2013) and rice against sheath blight disease (Sood et al. 2013), indicating that the developmental and maturation processes of treated plants have not been disturbed by the BTH exposure at the optimal dose and concentration level in different host plants. Also, the global transcriptomic analyses of wheat (Li et al. 2020), strawberry (Landi et al. 2017), and banana (Cheng et al. 2018) plants under BTH treatment indicated up-regulated genes associated with immune response. There are also reports on the positive effect of BTH treatment of tomato plants against *Botrytis cinerea* (Achuo et al. 2004) and activation of basal response to tomato spotted wilt virus and citrus exocortis viroid (López-Gresa et al. 2016, 2019). So far, the performed research has tended to focus on BTH as a SAR activator, when a positive relationship between marker genes of SA and JA (both increased after inducer treatment) was observed possibly suggesting the synergistic mode of action of these phytohormones in defence induction (Frackowiak et al. 2019) n. Although all the mentioned studies provide general information about the decrease in the accumulation level of those pathogens, the detailed mode of BTH action in tomato plants is still poorly understood.

Previously, we showed that the BTH and its ionic derivatives enhanced the plant immune response in *Nicotiana tabacum* - tomato mosaic virus (ToMV) pathosystem. Plants activated general immune response a few hours after BTH application and have shown up-regulated expression of SAR and ISR marker genes, including *PR-1b* and *PR-2, LIPOXYGENASE, ETHYLENE RECEPTOR* (Frackowiak et al. 2019). Moreover, in tobacco plants treated with the BTH either, before or after ToMV inoculation, the accumulation of viral RNA in systemic leaves was significantly reduced, indicating the priming effect of tested inducers on the analysed host plants (Frackowiak et al. 2019). We have also observed that the priming effects in BTH-treated tobacco plants were maintained up to a month after its application to soil.

This study aimed to comprehensively reveal the BTH mode of action in tomato plants, which has not been analysed before. Additionally, to determine the strength of immune priming to a viral pathogen, BTH-treated and non-treated plants were infected with ToMV. To decipher the overall BTH-induced response of the plant, the data obtained from transcriptomic and proteomic analyses were combined and enriched with the results of metabolomic analyses. We found out that the response of tomato plants to BTH treatment was related to changes in the expression level of *WRKY* transcription factors, *ARGONAUTE 2A (AGO2A)*, genes involved in thiamine and glutathione metabolism (with *GLUTHATHIONE S-TRANSFERASE* (*GST)* gene expression), cell wall reorganization, detoxification processes and differences in the amount of three phytohormones: ABA, JA-Ile, and Aux.

## 2. Materials and Methods

### 2.1. Tomato plants, chemical treatment with selected inducers and virus inoculation

The experimental model consisted of a host plant (*S. lycopersicum* cv. Betalux), a virus (SL-1 isolate of ToMV), and a resistance inducer - the Benzo(1,2,3)-Thiadiazole-7-Carbothioic Acid S-Methyl Ester (BTH). The plants were maintained in greenhouse conditions with a 16-hour day/ 8-hour night cycle in 26 °C day/23 °C night.

BTH was synthesized at the Poznań Science and Technology Park, AMU Foundation (Poland). It was used for watering six- to eight-week-old plants (100 mL) in the concentration of 10 mg/L. Initially, a group of twelve plants was treated with water and another twelve plants were treated with BTH. Next, water-treated and inducer-treated plants were collected 1-day post BTH treatment (1dpt). After one week, 4 plants previously water- or BTH-treated, were inoculated with the purified virus inoculum (at a concentration of 7.2×10^−4^ ng per leaf) by rubbing carborundum-dusted leaves. The second sampling of the plant material was performed at 8 dpt. Four replicates were performed for each analysed experimental condition. All collected samples were frozen in liquid nitrogen and stored at -80 °C. Portions of the collected plants from 1 and 8 dpt were freeze-dried using a laboratory freeze dryer - Alpha 1-4 L.S.C. basic (Martin Christ, Germany) and prepared for targeted hormonal analysis.

### 2.2. Isolation of total RNA and cDNA synthesis

For the RNA sequencing (RNA-Seq), the total RNA was isolated using the acidic phenol-chloroform method (Molnár et al. 2005) with some modifications. The isolated RNA was suspended in 30-50 μL of RNase-free water. Next, RNA was DNase-treated (Thermo Scientific, USA), and the purified RNA (between 3-5 μg) was stored at - 80 °C until use. The quality and concentration of the prepared RNA were assessed using a 2100 Agilent Bioanalyzer (Agilent, USA).

For validation of the obtained results, the total RNA was isolated using the RNeasy Plant Mini Kit (Qiagen, Germany). The purified RNA (2 μg) was used for cDNA synthesis using the RevertAid RT Reverse Transcription Kit (Thermo Scientific, USA) in the presence of 100 μM random hexamers (Thermo Scientific, USA).

### 2.3. The RNA-Seq and data analysis

The purified total RNA samples from all 24 plants were used for cDNA libraries synthesis. The libraries and RNA-seq were prepared by MGI Tech Co., Ltd. (China) as described before (Zhu et al. 2018). The quality check of the obtained raw data reads and adapter trimming was performed by Genomed S.A. (Poland). The filtered clean reads from each sample were aligned to the reference tomato genome of *S. lycopersicum* SL3.0 from the International Tomato Genome Sequencing Project (ftp://ftp.solgenomics.net/tomato_genome/assembly/build_3.00/) with annotation file ITAG3.2 (ftp://ftp.solgenomics.net/tomato_genome/annotation/ITAG3.2_release/). Mapping and subsequent analyses were made using the OmicsBox software (OmicsBox - Bioinformatics made easy. BioBam Bioinformatics (Version 1.2.4). March 3, 2019. www.biobam.com/omicsbox) (Dobin et al. 2013; Anders et al. 2015; Anders 2018). The count table obtained after the mapping process was filtered using the edgeR packet in the R environment (Version 3.6.2) (Robinson et al. 2009).

The differential gene expression analysis was carried out through a comparison between the treated and control groups from each time point of the experiment using OmicsBox (Robinson et al. 2009). The number of clean reads that mapped to each annotated transcript after RNA-Seq analysis was calculated using the HTSeq algorithm followed by the TMM (Trimmed Mean of M values) normalization method. The GLM (Quasi Likelihood-F test) statistical test was used in further data analysis. The DEGs were filtered by *p-Value* ≤ 0.05 and log2 fold change -1 ≥ (FC) ≥ 1.

### 2.4. Proteomic Analysis

For protein isolation, 0.2-0.4 g of plant material was homogenized using liquid nitrogen and in the presence of the extraction buffer (0.1M Tris-HCl, pH 6.8; 2mM EDTA-Na_2_; 20mM DTT and 2mM PSMF with 1% SDS) followed by centrifugation (15,000 ×g, 3 min, 4°C) to remove plant debris. Then 100% chilled acetone was added to the supernatant at the ratio (5:1) and left overnight at -20°C. The precipitate was centrifuged (5,000 ×g, 5 min, 4°C) and the resultant pellet was rinsed three times with 80% chilled acetone. The precipitate was then dried using a Vacuum Concentrator (Heraeus Instruments, Denmark) and stored at -80 °C until used.

A detailed description of the subsequent stages of protein analysis is presented in (Supplementary File 1A). To prepare a list of statistically significant DEPs, the results were filtered by using *p-Value* < 0.05 and log2FC ≤ 0.75 (decrease in abundance) or log2FC ≥ 0.75 (increase in abundance) as the threshold.

### 2.5. Functional analysis of DEGs and DEPs

The annotation file of *S. lycopersicum* genes was downloaded from the BioMart Ensembl database (https://www.ensembl.org/biomart/martview/e0215cf00bebb5d163991ca7859c5e80). To obtain functional annotation and identify putative biological pathways of statistically significant DEGs and DEPs separately, the NCBI Gene Ontology (GO) (Harris et al. 2004) and the Kyoto Encyclopaedia of Genes and Genomes (KEGG) (Kanehisa and Goto 2000) databases were used, all performed in the OmicsBox software. For functional analysis of DEPs, the protein ID was mapped to the Ensembl Genome Transcript IDs using the UniProt mapping service. The *p-Value* < 0.05 was used as a threshold to define significantly enriched GO terms (Fisher’s exact test) and KEGG pathways. To determine pathways enriched with transcriptomic and proteomic data, the combined file of DEGs and DEPs for each experimental condition was generated (including common terms). The OmicsBox software and R software with GOplot (Walter et al. 2015), ggplot2 (https://www.rdocumentation.org/packages/ggplot2/versions/3.3.3) and heatmap (https://github.com/raivokolde/pheatmap) packages were used to specify the enriched GO terms and to visualize the results of GO and KEGG analysis. The STRING application in the Cytoscape software was used (Doncheva et al. 2019) to determine the most statistically significant KEGG pathways common for all experimental conditions. To perform this analysis, all of the statistically significant DEGs and DEPs have been converted to STRING ID using the g:Profiler browser tool (g:Converter) (https://biit.cs.ut.ee/gprofiler/convert). Results were collected and presented using PowerPoint software.

### 2.6. Targeted hormonal analysis - liquid chromatography coupled to ESI mass spectrometry

The hormones were analysed as described by Sanchez-Bel et al. (2018) (Sánchez-Bel et al. 2018). A detailed description of the subsequent stages of targeted hormonal analysis is presented in the supplements (Supplementary File 1B). The Student’s t-test and Wilcoxon test were conducted in an R environment to examine significant differences between BTH- and water-treated samples. The *p-Value* was calculated for each experiment (with marking values cut-off <0.05).

### 2.7. The RT-qPCR validation

Real-time quantitative PCR (RT-qPCR) was performed with gene-specific primers designed using the Primer3 (v. 0.4.0) software. Each RT-qPCR reaction (10 μl) included 2 × iTaq™ Universal SYBR®Green Supermix (Bio-Rad, CA, USA), 500 nM forward and reverse primers (Supplementary File 1C), and 1 μL of cDNA. PCR thermal cycling was performed as described previously (Wrzesińska et al. 2018).

Gene expression studies were performed for four biological replicates, and for each biological replicate, three technical replicates were analysed. Relative gene expression was calculated using the GenEx software (MultiD Analyses AB, Sweden) and REST software (Pfaffl et al. 2002). An independent non-parametric Mann-Whitney test was conducted to test for significant differences between inducer (BTH)- and water-treated samples in all experiments. The *p-Value* was calculated for each experiment (with marking values cut-off <0.05 (*)).

## 3. Results

### 3.1 The general transcriptomic (RNAseq) and proteomic data analysis

Plants (leaves) were harvested at two-time points, on day 1 and day 8 post the BTH treatment (dpt). Additionally, seven days after the BTH or water (control/mock) treatment, the plants were inoculated with ToMV and their leaves were harvested one day after virus inoculation, to evaluate the impact of BTH on the virus-infected plant (8dpt/1dpi) (Fig. 1).

**Fig. 1.**
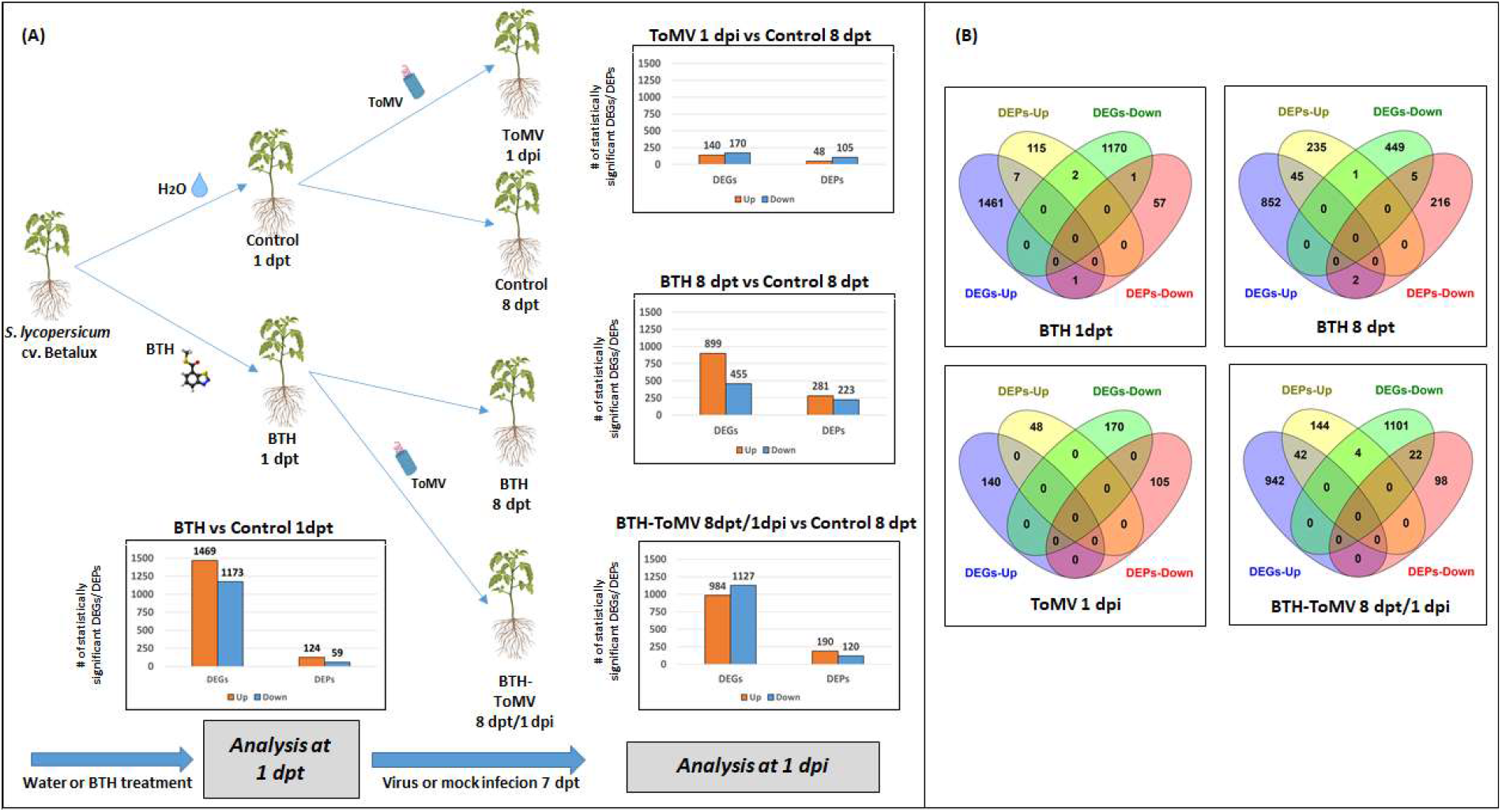
A) The scheme presents the experimental setup used in the study. *Solanum lycopersicum* plants were treated with BTH or water (control). One day post-treatment (1dpt) the first plants were harvested from the control 1 dpt and BTH-treated (BTH 1dpt) plants. One week after BTH application, the groups of water- and BTH-treated plants were inoculated with ToMV. On the next day, 1-day post-infection (1 dpi), plant material was collected from control 8 dpt, non-infected and BTH-treated plants (BTH 8 dpt), BTH-treated-ToMV-infected plants (BTH-ToMV), and water-treated-ToMV-infected plants (ToMV). Graphs showing the number of statistically significant differentially expressed genes - DEGs (two left bar plots on each graph) and differently expressed proteins DEPs (two right bar plots on each graph). The orange bar plots present up-regulated DEGs/DEPs opposite to the blue bar plots presenting down-regulated DEGs/DEPs; B). The Venn diagram represents the overlapping up- and down-regulated DEGs and DEPs numbers in each experimental condition [Created with BioRender.com].

In general, 21,834 annotated transcripts for all experimental conditions were obtained (Supplementary Table 1) with the highest number of statistically significant differentially expressed genes (DEGs) identified in BTH-treated plants at 1 dpt (1,469 up-regulated, 1,173 down-regulated) and the lowest number of DEGs identified for ToMV-infected plants (1 dpi) (140 up- and 170 down-regulated) compared to control plants at each time point (Fig. 1). The list of all DEGs is presented in Supplementary Table 2.

The proteomic analysis resulted in the indication of 3,459; 3,562; 3,559; and 3,460 annotated proteins for BTH-treated plants: BTH 1 dpt, BTH 8 dpt, BTH**-**treated-ToMV-infected 8 dpt/ 1 dpi, and ToMV-infected 1 dpi plants, respectively (Supplementary Table 1B). The filtered differentially expressed proteins (DEPs) data (*p-Value* < 0.05) showed the highest number of DEPs identified for BTH-treated plants 1 dpt (281 increased and 223 decreased), and the lowest number of DEPs identified for ToMV-infected plants (1 dpi) (48 increased and 105 decreased) (Fig. 1). The list of all DEPs is presented in Supplementary Table 2.

### 3.2 The main biological processes and KEGG pathways involved in tomato response to BTH

To evaluate the impact of BTH treatment on tomato plants, the functional analysis of transcriptomic and proteomic datasets was performed. The Gene Ontology (GO) terms were reduced to the most specific ones, followed by the selection of the 15 most enriched biological processes for each type of experimental condition (Supplementary Table 3). The KEGG pathway analysis results were also reduced to the 20 most enriched in plants at the early (1dpt) and late (8 dpt) phase of response to BTH. This analysis was also performed for ToMV-infected and non-infected plants (Supplementary Table 4).

At the early stage of tomato plants’ response to BTH treatment (1 dpt), the majority of up-regulated processes at the transcriptional level were associated with the detoxification process, photosynthesis, glutathione metabolic process, defence induction, ribosomal assembly process, and auxin (Aux)-mediated pathway related to root development. The regulation of the root organization process was also presented in the up-regulation of DEGs related to root development and proteins connected to lateral root formation and auxin polar transport. Also, 5 up-regulated DEPs connected to the response to the ABA process were noted. Both ABA and Aux may play a key role in the first stage of tomato response to BTH application. Interestingly, the transcripts associated with the protein phosphorylation process and positive regulation of transcription by RNA polymerase II were mostly down-regulated (71 and 18 of tested transcripts, respectively) (Supplementary Table 3).

In contrast to the early response, in BTH-treated tomato plants 8 dpt, the up-regulations of DEGs and DEPs associated with the protein phosphorylation process and down-regulation of the photosynthesis process were observed. Moreover, we noted changes in the cell wall reorganization process, namely up-regulation of cell wall macromolecule catabolic process (transcriptomic data) and chitin catabolic process (transcriptomic and proteomic results), together with down-regulation of cell wall organization and biogenesis processes, xyloglucan metabolic process, and cellulose catabolic process (transcriptomic datasets). Likewise as in the early stage, in the late phase of tomato response to BTH application the up-regulation of glutathione metabolic process, detoxification process, and defence response to fungus and bacterium were noted at transcriptomic and proteomic levels. Furthermore, at the transcriptome level, the up-regulation of SAR and plant-type hypersensitive response (HR) were noted (Supplementary Table 3).

After viral infection in mock- and BTH-pre-treated plants, we detected an increased number of DEGs and DEPs associated with the response to oxidative burst (response to oxidative stress, hydrogen peroxide catabolic process, cellular oxidant detoxification) and a reduced number of transcripts and proteins linked to photosynthesis process (light-harvesting in photosystem I, electron transport in photosystem II, photorespiration, and chlorophyll biosynthetic process) than in BTH-treated tomato plants 1 dpt. Noteworthy, in the water-treated-ToMV-infected tomato plants we observed up-regulation of the ABA catabolic process (transcriptomic data) together with down-regulation of thiamine (vitamin B1) biosynthetic process (proteomic data), which play a role in plant developmental process and defence induction in plant response to ToMV infection (Supplementary Table 3).

The described above results were also confirmed by KEGG pathway analysis. In all experimental conditions, the commonly represented pathways were starch and sucrose metabolism (mostly down-regulated at transcriptome level of BTH-treated plants both for 1 dpt and 8 dpt, and up-regulated in both transcriptome and proteome levels of BTH-ToMV-treated plants 1 dpi) and purine metabolism (highly up-regulated in BTH-treated plants 1 dpt and BTH-treated-ToMV-infected plants 8dpt/1dpi). Interestingly, the thiamine metabolism process was greatly up-regulated at transcriptome level in BTH-treated plants 1 dpt, at proteome level in BTH-treated plants 8 dpt, and at both transcriptome and proteome levels in BTH-ToMV-treated plants 1 dpi. At the early response of tomato plants to BTH treatment and after virus infection, the up-regulation of drug xenobiotic biodegradation and metabolism were noted, which was associated with detoxification and regulation of oxidative burst processes. Changes in the carbon fixation in photosynthetic organisms pathways were up-regulated in BTH-treated tomato plants 1 dpt and down-regulated in BTH-treated mock- (8 dpt) or ToMV-infected (1 dpi) plants (Supplementary Table 4).

### 3.3 The transcriptomic and proteomic datasets comparison

All DEGs and DEPs records were unified and the combined data were subjected to functional analysis. The plant processes and KEGG pathways, statistically most significant, were defined using the GOplot package in the R environment and Cytoscape software with String Enrichment application. Functional analysis of tested DEGs and DEPs showed enrichment of many processes and signalling pathways in BTH-treated plants, including primary metabolism, transcription and translation, cell wall organization, phosphorylation, photosynthesis, detoxification, response to phytohormone, and defence response (Fig. 2).

**Fig. 2.**
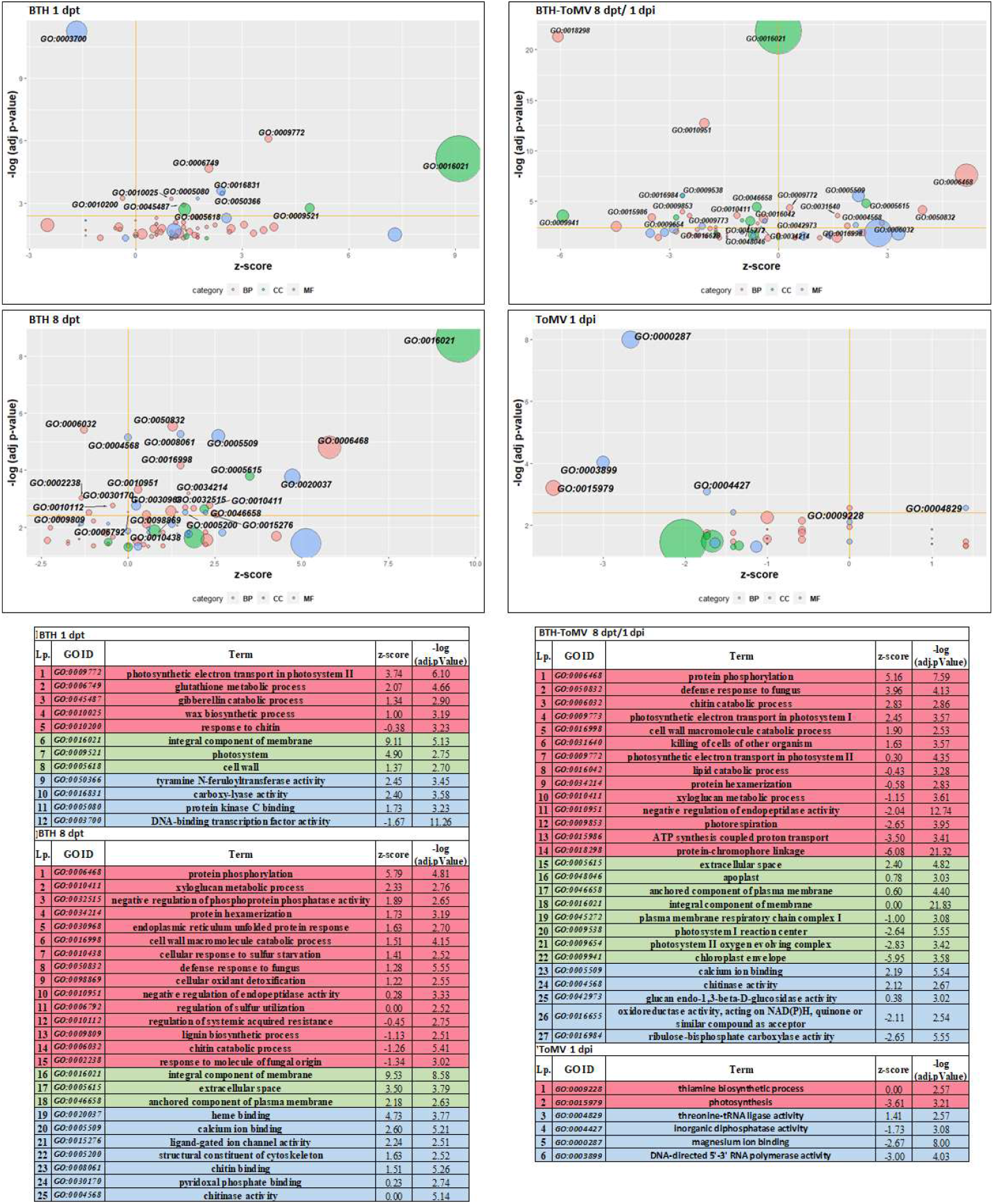
Bubble plots generated with GOplot R package showing statistically significant enriched GO terms for each type of experimental conditions after analysis of combined DEGs and DEPs data. Plots were divided into four groups dependent on experimental variants (BTH-treated plants 1dpt – upper left, BTH-treated plants 1 dpi – down left, BTH-treated-ToMV-infected plants 1 dpi – upper right, and water – treated-ToMV-infected plants 1 dpi– down right). The size of bubbles is proportional to the number of DEGs and DEPs unified IDs assigned to each GO term. The x-axis represents the z-score and the y-axis represents the negative logarithm of the adjusted *p-Value* calculated in the R package. The threshold was set to 2.5 of –log(adj *p-Value*). Each category of GO term is represented by a different colour: Biological Process – red; Cellular Component – green and Molecular Function – blue.

Similarly, as noted above, increased levels of the DEGs and DEPs associated with the photosynthesis, glutathione metabolic process, and gibberellin catabolic process were observed for BTH-treated plants 1 dpt (Fig. 2).

At the late phase of response, in BTH-treated plants 8 dpt, the reorganization of the cell wall was also noted (by up-regulation of xyloglucan metabolic process, cell wall macromolecule catabolic process, chitin-binding, and chitinase activity, and down-regulation of lignin biosynthetic process and chitin catabolic process). Comparison of the transcriptomic and proteomic datasets obtained for BTH-treated plants 8 dpt showed that the protein phosphorylation and defence response processes were much up-regulated (Fig. 2).

In the BTH-treated-ToMV-infected plants 8 dpt/1 dpi, the up-regulation of processes concerning protein phosphorylation, cell wall reorganization process, and defence response were observed. Interestingly, the photosynthetic electron transport processes in photosystems I and II were up-regulated, while the photosystem reaction centre I, photosystem II oxygen-evolving complex, and chloroplast envelope components were highly down-regulated (Fig. 2).

The KEGG analysis showed significant changes in pathways associated with thiamine metabolism (up-regulation in BTH-treated plants 1 and 8 dpt, and down-regulation in BTH-ToMV-treated tomato plants) and up-regulation in phenylpropanoid biosynthesis and phenylalanine metabolism in all experimental variants of BTH-treated plants. In healthy BTH-treated plants (1 and 8 dpt) a slight increase in stilbenoid, diarylheptanoid, and gingerol biosynthesis was observed. At the late response to BTH treatment of tomato plants, an increase in ubiquinone and another terpenoid quinone biosynthesis was indicated together with a decrease in glyoxylate and dicarboxylate metabolism (both healthy and ToMV-infected plants). There was also a difference between healthy BTH-treated plants 8 dpt and BTH-ToMV-treated plants in purine metabolism, namely this process increased in healthy plants, while decreased after virus infection (Fig. 3). Drug metabolism, glutathione metabolism, and flavonoid biosynthesis were noticeably up-regulated in BTH-treated plants 1 dpt (Fig. 3). What is striking, all processes presented for water-treated-ToMV-infected plants were down-regulated (Fig. 3).

**Fig. 3.**
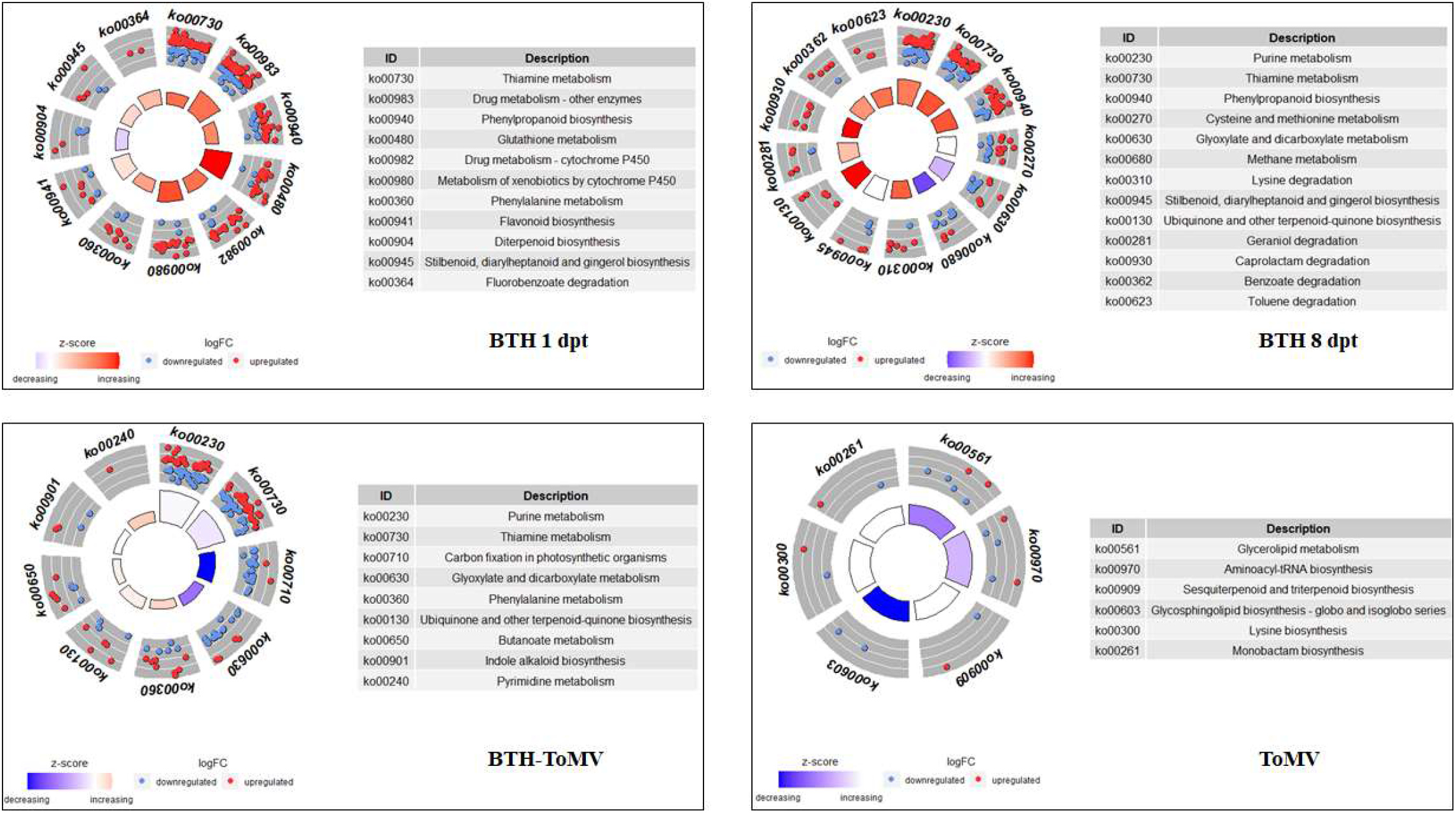
Statistically significant KEGG pathways, previously analysed in the OmicsBox software. The visualisation was generated in the R environment (GO plot R package). Each dot represents one DEG/DEP (red dot – up-regulated or blue dot – down-regulated). The outer radius of the circle contains the pathway ID (explained in each table), while the internal radius represents the statistical significance (trapezoid size) and the direction of regulation of the presented pathway (based on the Z-score).

The significant up-regulation of pathways associated with detoxification and oxidative stress regulation, defence induction (plant-pathogen interaction), and hormone signalling regulation were also noted in the BTH-treated tomato plants. Also, the ribosome pathway was much up-regulated in the BTH-treated plants at 1 dpt (46 of all associated DEGs/DEPs) and protein kinase activity process and phenylpropanoid biosynthesis pathways (with phenylalanine metabolism pathway) were mostly up-regulated in the BTH-treated plants 8 dpi (Fig. 4).

**Fig. 4.**
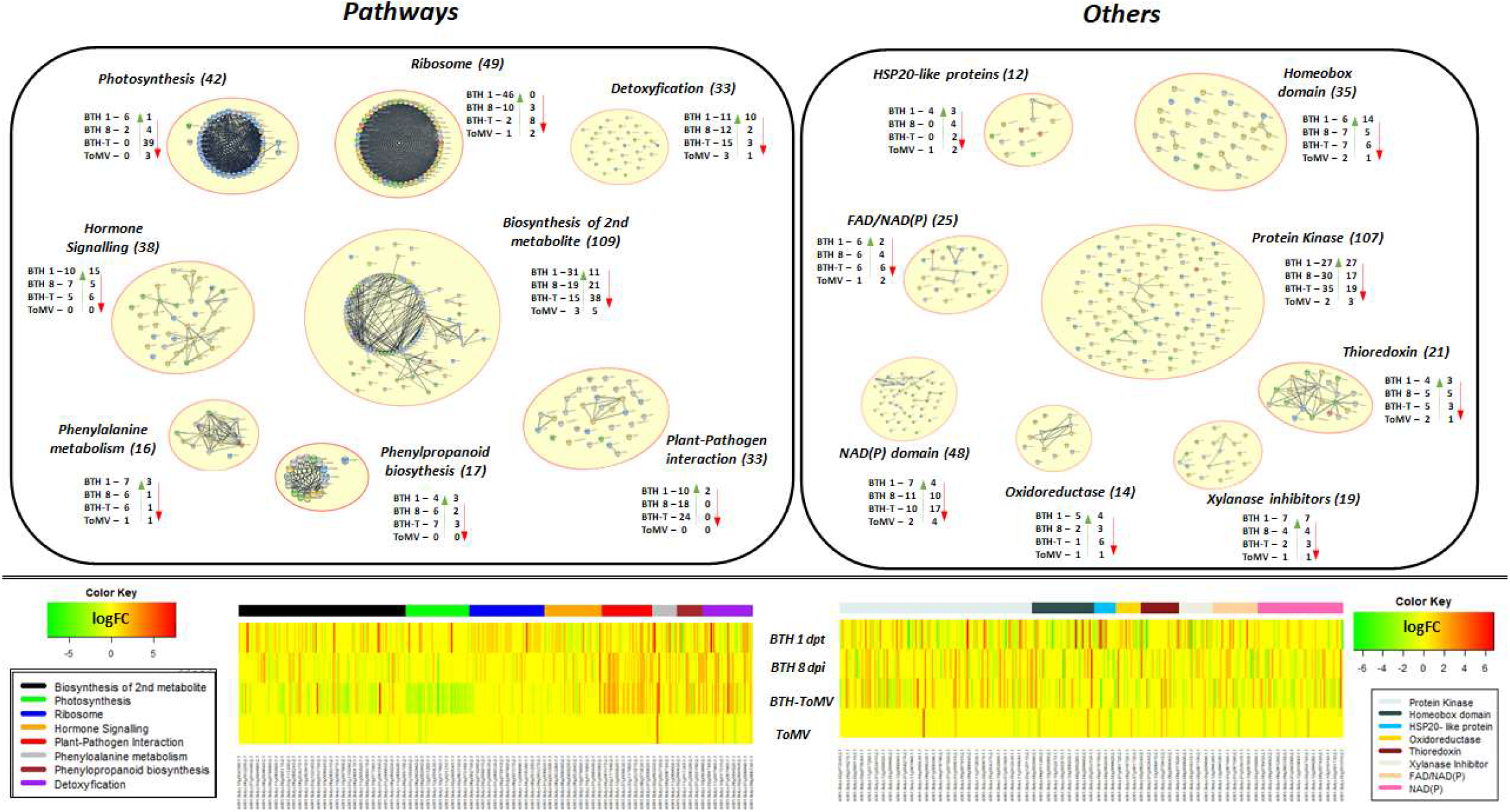
Maps of selected pathways, enzymes, DEGs/DEPs associated with tomato plants’ response to BTH treatment (1 dpt and 8 dpt) and/or virus-infected (treated with BTH [BTH-ToMV] or with water [ToMV]) received from Cytoscape application. The number of found genes associated with each term is presented in brackets. Nodes have been grouped by experimental condition, where the green arrows show the number of up-regulated genes and the red arrows show the number of down-regulated genes. The bottom part presents heatmaps generated from the logFC values of each process/enzyme/protein presented above from each experimental condition. Each colour bar represents each term that is shown above and indicates the range of genes observed for a given term on heatmaps.

### 3.4. The effect of BTH treatment on changes in the levels of phytohormones in tomato plants

To determine the impact of BTH on the levels of phytohormones, seven different hormones were selected: ABA, SA, three associated with the JA pathway: JA-Ile, JA, and cis-(+)-12-oxo-phytodienoic acid (OPDA), and two connected with the auxin pathway: indole-3-acetic acid (IAA) and I3CA (Table 1).

**Table 1.**
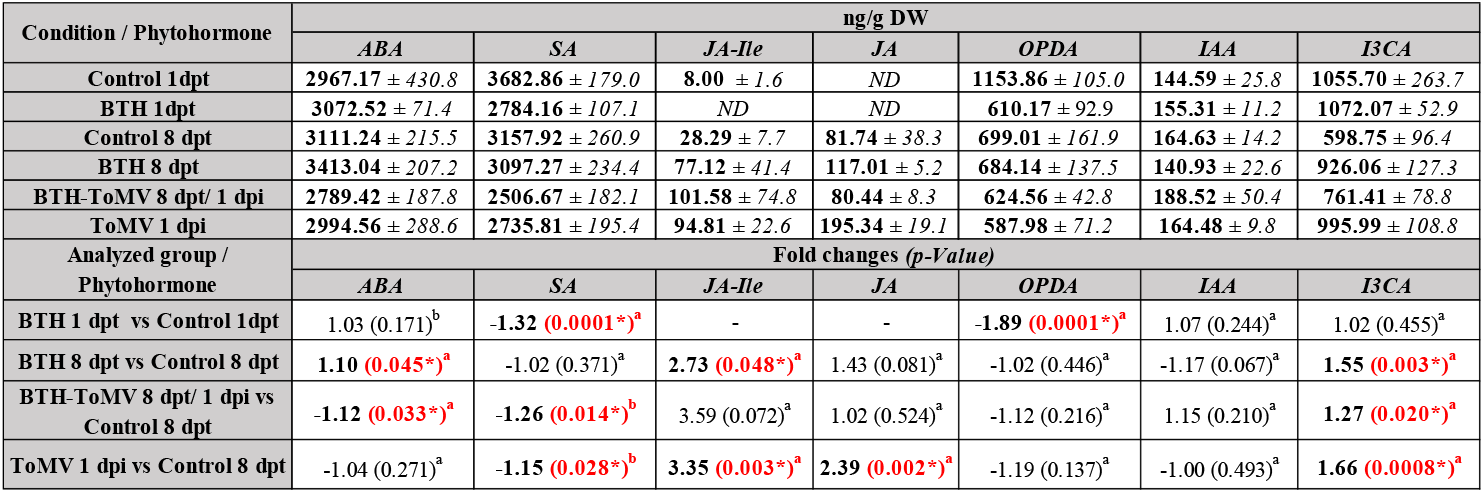
Results of differences in the amounts of synthesized phytohormones acquired from the analysed plants calculated on a dry basis. The first value represents mean results from at least 3 plants from each experimental condition in ng/g DW (dry weight). The second value (after ±) is recalculated SD (standard deviation) from each result. ABA – Abscisic Acid; SA – Salicylic Acid; JA-Ile – Jasmonic Acid-Isoleucine; JA – Jasmonic Acid; OPDA - cis-(+)-12-oxo-phytodienoic acid; IAA - Indole-3-acetic acid; I3CA - Indole- 3-carboxylic Acid; ND – not detected. Statistically significant results were presented in the bottom panel of the table in red (calculated using Student t-Test (a) or Wilcoxon Test pointed (b)).

| Condition / Phytohormone | ng/g DW |  |  |  |  |  |  |
| --- | --- | --- | --- | --- | --- | --- | --- |
|  | ABA | SA | JA-Ile | JA | OPDA | IAA | I3CA |
| Control 1dpt | 2967.17 ± 430.8 | 3682.86 ± 179.0 | 8.00 ± 1.6 | ND | 1153.86 ± 105.0 | 144.59 ± 25.8 | 1055.70 ± 263.7 |
| BTH 1dpt | 3072.52 ± 71.4 | 2784.16 ± 107.1 | ND | ND | 610.17 ± 92.9 | 155.31 ± 11.2 | 1072.07 ± 52.9 |
| Control 8 dpt | 3111.24 ± 215.5 | 3157.92 ± 260.9 | 28.29 ± 7.7 | 81.74 ± 38.3 | 699.01 ± 161.9 | 164.63 ± 14.2 | 598.75 ± 96.4 |
| BTH 8 dpt | 3413.04 ± 207.2 | 3097.27 ± 234.4 | 77.12 ± 41.4 | 117.01 ± 5.2 | 684.14 ± 137.5 | 140.93 ± 22.6 | 926.06 ± 127.3 |
| BTH-ToMV 8 dpt/ 1 dpi | 2789.42 ± 187.8 | 2506.67 ± 182.1 | 101.58 ± 74.8 | 80.44 ± 8.3 | 624.56 ± 42.8 | 188.52 ± 50.4 | 761.41 ± 78.8 |
| ToMV 1 dpi | 2994.56 ± 288.6 | 2735.81 ± 195.4 | 94.81 ± 22.6 | 195.34 ± 19.1 | 587.98 ± 71.2 | 164.48 ± 9.8 | 995.99 ± 108.8 |
| Analyzed group / Phytohormone | Fold changes (*p*-Value) |  |  |  |  |  |  |
|  | ABA | SA | JA-Ile | JA | OPDA | IAA | I3CA |
| BTH 1 dpt vs Control 1dpt | 1.03 (0.171)<sup>b</sup> | -1.32 (0.0001*)<sup>a</sup> | - | - | -1.89 (0.0001*)<sup>a</sup> | 1.07 (0.244)<sup>a</sup> | 1.02 (0.455)<sup>a</sup> |
| BTH 8 dpt vs Control 8 dpt | 1.10 (0.045*)<sup>a</sup> | -1.02 (0.371)<sup>a</sup> | 2.73 (0.048*)<sup>a</sup> | 1.43 (0.081)<sup>a</sup> | -1.02 (0.446)<sup>a</sup> | -1.17 (0.067)<sup>a</sup> | 1.55 (0.003*)<sup>a</sup> |
| BTH-ToMV 8 dpt/ 1 dpi vs Control 8 dpt | -1.12 (0.033*)<sup>a</sup> | -1.26 (0.014*)<sup>b</sup> | 3.59 (0.072)<sup>a</sup> | 1.02 (0.524)<sup>a</sup> | -1.12 (0.216)<sup>a</sup> | 1.15 (0.210)<sup>a</sup> | 1.27 (0.020*)<sup>a</sup> |
| ToMV 1 dpi vs Control 8 dpt | -1.04 (0.271)<sup>a</sup> | -1.15 (0.028*)<sup>b</sup> | 3.35 (0.003*)<sup>a</sup> | 2.39 (0.002*)<sup>a</sup> | -1.19 (0.137)<sup>a</sup> | -1.00 (0.493)<sup>a</sup> | 1.66 (0.0008*)<sup>a</sup> |

In BTH-treated plants 1 dpt, a slight increase in the amount of IAA and a decrease in the amounts of ABA and I3CA were observed, while the SA and the OPDA accumulation levels were lower than in control plants (1 dpt) 1.32 times and 1.89 times, respectively. In BTH-treated, non-infected plants (8 dpt) statistically significant changes were observed in the levels of phytohormone ABA (1.10 times higher than in control plants), and an increase was observed in I3CA (1.55 times) relative to control plants. Also, a high increase in the level of JA-Ile (2.73 times) was noted for tomato plants BTH-treated 8 dpt.

The ToMV infection affected the abundance of the tested phytohormones in both BTH-treated and water-treated tomatoes. In the infected plants, in comparison to the control plants, BTH caused slight changes in the amount of ABA (1.12 times lower), SA (1.26 times lower), and I3CA (1.27 times higher). Interestingly, the level of JA-Ile was 3.59 times higher than in control plants (statistically insignificant). In virus-infected plants, treated only with water, an increase in JA-Ile (3.35 times), JA (2.39 times), and I3CA (1.66 times), and a decrease in SA (1.15 times), relative to controls, was observed.

### 3.5. Validation of the obtained results

To validate the obtained results, 18 genes associated with 12 processes (namely: “signalling”, “gene expression”, “defence response”, “response to stress”, “detoxification”, “response to hormone” (with a detailed response to Aux and JA), “glutathione metabolic process”, “water-soluble vitamin biosynthetic process”, “programmed cell death” and “response to wounding”) were validated by RT-qPCR (Fig. 5 and Supplementary File 1c).

**Fig. 5.**
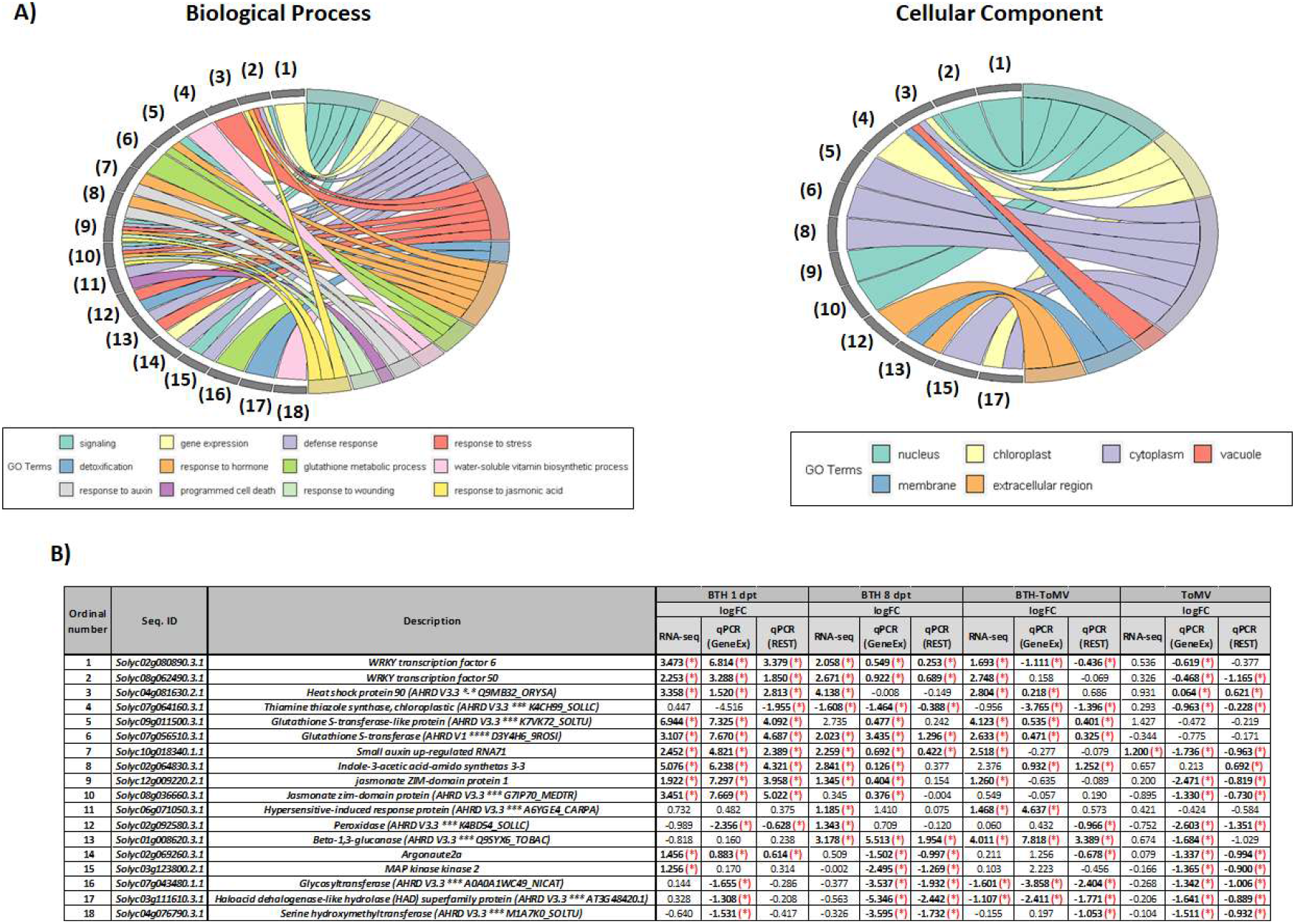
A) The assignment of validated genes to Biological Process and Cellular Component; B) Table presents logFC results for each validated gene divided into results obtained from RNA-seq and RT-qPCR (two different calculation software: GeneEx ver 6 and REST) analysis. The statistically significant results were marked with a red star (p-Value < 0.05, calculated using the Mann-Whitney test).

The highest number of statistically significant results was identified for BTH-treated plants 1 dpt, opposite to BTH-treated and ToMV infected tomato plants (Fig. 5b). The highest changes in expression profile were observed for *WRKY TRANSCRIPTION FACTOR 6, WRKY TRANSCRIPTION FACTOR 50, GST* (two transcripts), *SMALL AUXIN UP-REGULATED RNA71, INDOLE-3-ACETIC ACID-AMIDO SYNTHETASE 3-3, JASMONATE ZIM-DOMAIN PROTEIN 1* (*Solyc12g009220.2.1*), and *JASMONATE ZIM-DOMAIN PROTEIN 11* (*Solyc08g036660.3.1*), all of which were up-regulated in BTH-treated tomato plants 1 dpt and 8 dpt. Down-regulation in *PEROXIDASE* expression was observed for BTH-treated tomato plants 1 dpt and ToMV-infected water-treated tomato plants. *GLYCOSYLTRANSFERASE* and *HOLOACID DEHALOGENASE-LIKE HYDROLASE (HAD) SUPERFAMILY PROTEIN* genes were down-regulated in BTH-treated tomato plants 8 dpt (both un- and infected with ToMV). In turn, *PR2* gene expression was highly up-regulated in the same experimental variants. The expression of *THIAMINE THIAZOLE SYNTHASE* gene was decreased in all experimental conditions (highly declined in BTH-treated tomato plants 1 dpt and BTH-ToMV-treated tomato plants), and increased expression of *AGO2A* gene in BTH- treated plants 1 dpt (as opposed to water-treated-ToMV-infected tomato plants, where expression of this gene involved in post-transcriptional gene silencing (PTGS) was decreased) (Fig. 5b).

## 4. Discussion

### Changes in differentially expressed genes (DEGs) and proteins (DEPs) upon BTH treatment

BTH is known as a synthetic plant protection agent that mimics SA properties and enhances the plant immune system before the appearance of a pathogen, which has been confirmed for many plant species, among others, rice (Sood et al. 2013), strawberry (Landi et al. 2017), banana (Cheng et al. 2018) and potatoes (Brouwer et al. 2020). In the present study, BTH-induced resistance of tomato plants from in-depth transcriptomic, proteomic, and targeted metabolomics analyses were described. For the first time, early and later phases of *S. lycopersicum* response to BTH were analysed in detail, including the effect on the initial stage of viral infection. BTH application to tomato plants resulted in a high increase in the DEGs number at 1 dpt, while in BTH-treated plants 8 dpt fewer DEGs have been identified. The contrary state was observed for DEPs, where an increase of a higher number of DEPs was noticed at 8 dpt rather than 1 dpt.

### BTH treatment causes up-regulation of WRKY transcription factors genes in tomato plants

Transcription factors (TFs) play one of the most important roles in the cell machinery, including WRKY family transcription factors, which were identified amongst the most up-regulated TFs in plants upon BTH treatment. Li et al. (2020) have highlighted the initiation of WRKY family gene expression in the plants after BTH treatment, postulating their indirect effect on immunity induction through the synthesis of genes from the PR family in an NPR1-independent way (Li et al. 2004, Li et al. 2020). In our study, a higher expression of *WRKY* genes was observed, of which 3 are related to the developmental process (*WRKY6/35/75*), while 5 are associated with the plant’s defence response (*WRKY 33/50/51/80*/*81*) (Rushton et al. 2010). The up-regulation of two genes belonging to the WRKY family, namely: *WRKY6* (*Solyc02g080890*) and *WRKY50* (*Solyc08g062490*), was presented in BTH-treated tomato at each analysed time point (with the highest expression change at 1 dpt). Those two genes are supposed to be activated by BTH application and may have a crucial role in maintaining the metabolic balance of tomato plants by regulating the transcription process in the first phase of tomato response to BTH treatment.

### Impact of BTH treatment on post-transcriptional gene silencing process in tomato

The plant defence response is also related to the process of gene silencing, including PTGS (Vaucheret et al. 2001). In *Arabidopsis thaliana* pre-treated with BTH, the enhanced gene expression of the *AGO*2 gene was observed due to cucumber mosaic virus infection (CMV) (Ando et al. 2021). The AGO proteins, together with small RNAs, are involved in the formation of the RNA-induced silencing complex (RISC), the core element in the PTGS (Grene 2002). In this study, a significant increase in the expression level of *AGO2A* and *DICER-LIKE 2D* (*DCL2D*) (*Solyc11g008530*) genes in BTH-treated plants at 1 dpt was detected. In late response to BTH treatment of tomato plants, including virus-infected ones, the expression level of *AGO2A* gene was decreased, when an increase in the AGO2A protein level was observed. These findings may suggest that *AGO2* and *DCL2D* may be also involved in the defence response to ToMV infection in tomatoes.

### Regulation of immune response in BTH-treated tomato plants

Under pathogen infection in *Nicotiana benthamiana*, the GST attenuated the oxidative stress inside chloroplasts, enabling the synthesis of viral minus-strand of RNA (Budziszewska and Obrępalska-Stęplowska 2018). GST is associated with the glutathione metabolic process and is responsible for signalling processes in the plant, which was also observed in the presented results (e.g. the increased expression of *Solyc09g011500* and *Solyc07g056510* in all BTH-treated plants). However, a large number of GST transcripts were also associated with the response to hormones (mainly Aux) (e.g. mentioned before *Solyc09g011500*) and defence response (Supplementary Table 5).

The protein kinases play essential roles in the cellular signalling processes and control the mechanisms of protein modification and activation in plant-pathogen interactions (Kersten et al. 2009). The analysis of tomato plants’ response to BTH (1 dpt and 8 dpt) indicated a contribution of some of the analysed DEGs and DEPs in the phosphorylation process and kinase activity function (Fig. 5 and Supplementary Table 5). The increased expression of protein kinases genes (among others *MAP KINASE KINASE 2, Solyc03g123800*) and protein (MAP KINASE KINASE 1, *Solyc12g009020*) in virus-infected plants and their important role in the pathogenesis process have been described before (Wrzesińska et al. 2018, 2021), also for BTH-treated plants (Cheng et al. 2018). Such large changes in protein kinase levels probably contribute to maintaining the balance between the growth and the plant’s defensive functions, which is supported by the lack of visible phytotoxicity effects on BTH-treated plants.

At the early stage of tomato response to BTH, the increased expression of defence protein gene precursor, signal transduction protein (GST), and genes associated with hormone response (like *JAZ* genes *Solyc12g009220* and *Solyc08g036660*) was observed. Our previous research has revealed that BTH and its derivatives have a positive effect on defence induction not only related to SAR-type response but also cause changes in JA marker genes expression (like *JAZ* gene), which is associated with induced systemic resistance (ISR) (Frackowiak et al. 2019). A likely explanation is that both types of plant defence responses (SAR and ISR) are activated during BTH application to tomato and tobacco plants. In the late phase of tomato response to BTH treatment, the increased expression of *PR genes* (including the *PR2*) was also reported in tobacco plants (Frackowiak et al. 2019), and increased expression level of genes associated with SAR and programmed cell death (such as *Solyc06g071050, HYPERSENSITIVE-INDUCED RESPONSE PROTEIN*) was also noted in this study.

The response of tomato plants to BTH included changes in expression levels of DEGs and DEPs associated with the organization of the cell wall and its modification in all experimental conditions. The up-regulation of cell wall macromolecule catabolic process in BTH-treated tomato plants 8 dpt (both healthy and ToMV-infected plants) together with down-regulation of the lignin biosynthetic process in BTH-treated tomato plants 8 dpt were observed. Also decrease in the expression level of *GLYCOSYLOTRANSFERASE* (*Solyc07g043480*), which is involved in lignin biosynthesis pathways (Scheible and Pauly 2004), was confirmed for all experimental conditions of BTH-treated tomato plants. Together with the up-regulation of *PR2* gene expression, these findings indicate that the plant’s response is more complex and possibly explains why BTH can act on a wider spectrum of pathogens, viruses (Frackowiak et al. 2019), bacteria (Brisset et al. 2000), fungi (Araujo et al. 2015), and some pests (Inbar et al. 1998)).

### Changes in photosynthesis and vitamin B1 metabolism upon BTH treatment

Initial results suggest that there may be a connection between BTH application to tomato plants and changes in the photosynthesis process. Photosynthesis leads to the production of oxygen, energy (ATP), and carbohydrates, and the biosynthesis of many primary compounds, vitamins, phytohormones (SA, JA, ABA), and defence proteins. Chloroplasts participate in the immune response through the ROS/redox system (Grene 2002) and changes in cytosolic Ca^2+^ concentration (Lu and Yao 2018), but they are also associated with virus replication (Budziszewska and Obrępalska-Stęplowska 2018). At the early phase of tomato response to BTH, the photosynthesis-associated processes are highly up-regulated, while in the BTH-treated plants 8 dpt (both in virus-free and virus-infected plants), the photosynthesis was down-regulated. It may indicate that this process is crucial in the cell machinery stabilization after BTH action, which has been also observed for strawberries and bananas (Landi et al. 2017; Cheng et al. 2018). The up-regulation of the thiamine (vitamin B1) metabolism pathway, bringing a highly significant contribution to the plant’s defence response and development (Bocobza and Aharoni 2014; Dong et al. 2016), was observed in BTH-treated tomato plants 1 and 8 dpt. What is interesting, two genes involved in the thiamine metabolism pathway, showed the opposite direction of the expression level in BTH-treated plants. The first one, *HEAT SHOCK PROTEIN 90* (*Solyc04g081630*), is associated with plant response to biotic and abiotic stresses (including defence response to fungus and response to heat stress) (Xu et al. 2012). This gene was up-regulated in BTH-treated 1 dpt and BTH-treated-ToMV-infected plants, while the second gene, *THIAMINE THIAZOLE SYNTHASE* (*Solyc07g064160*), associated with thiamine biosynthesis, was highly down-regulated in all analysed BTH-treated plants. A former study confirmed the same situation when thiamine metabolism was increased with a reduction in its availability in plant stress conditions (Lee et al. 2007). Vitamin B1 acts as an enzymatic coenzyme in the maintenance of health and metabolism balance in plants (Fitzpatrick and Chapman 2020) and regulation of the response to abiotic (*via* ABA) and biotic stresses (Goyer 2010). It has been postulated that the increase in the expression of the genes associated with the thiamine metabolism pathway is correlated with the activity of SA (Ahn et al. 2005) and ABA (Rapala-Kozik et al. 2012) in rice, Arabidopsis, and cucumber. This is the first time that changes in the thiamine metabolism were found to correlate with BTH application.

### BTH application has an impact on levels of I3CA and JA-Ile

The phytohormones analysed in this study (ABA, SA, JA, and Aux) are associated, among others, with growth, signalling, transport, and immunity. Interestingly, the SA level was decreased at the early stage of tomato response to BTH application and in the plants infected with ToMV (both water- and BTH-treated). The decrease in the amount of SA in the BTH-treated plants both at 1 dpt and 8 dpt may be associated with the functional properties of the BTH molecule. The virus also has an impact on SA metabolism, both in water- and BTH-treated tomato plants, because a decrease in the amount of free SA in the plants was observed (higher in BTH-pre-treated tomato plants). Interestingly, an increase in the genes expression associated with the JA metabolic pathway in the BTH-treated plants (*Solyc12g009220* and *Solyc08g036660*) was detected and the amount of JA-Ile was also increased, with statistically significant results for the BTH-treated 8 dpt plants. JA-Ile is a JA derivative synthesized in the cytoplasm and together with the increase in *JAZ* genes expression, it regulates the plant immune response and developmental processes such as growth, seed germination, and root formation (Ghorbel et al. 2021). This observation confirms the results previously reported for *N. tabacum* – another plant from the Solanaceae family (Frackowiak et al. 2019), in which the expression levels of the genes related to the JA pathway also increased after the treatment with BTH or its derivatives. It seems that the application of BTH to plants influences a synergistic relation between SA- and JA-mediated pathways making the plant’s defence response probably more effective. In the late response to BTH treated plants 8 dpt, the amount of ABA and I3CA increased. For the BTH-treated-ToMV-infected tomato plants (8 dpt/1 dpi) an increase in the amounts of IAA and I3CA hormones was noted, while the level of ABA was decreased. Also, an up-regulation of gene expression associated with Aux/IAA (*Solyc10g018340*) and I3CA (*Solyc02g064830*), with the highest expression for BTH-treated plants 1 dpt was shown. IAA is important for plant root development and growth; it also plays a key role in the induction of *AUXIN RESPONSE FACTOR PROTEIN* gene expression and triggers the cellular immune response (Bari and Jones 2009). It has been reported that an increase in the amount of I3CA also induces ABA production (Gamir et al. 2018), which corresponds to callose deposition due to BABA treatment (Gamir et al. 2012). The data reported here appear to support the assumption that BTH influences the JA and I3CA biosynthesis pathways, but in general, it requires more studies to confirm that hypothesis.

### Conclusion

In conclusion, in this study, for the first time, a multi-omics analysis was carried out to decipher the tomato response to BTH administration in tomato plants. This study indicated that tomato plants’ response at the early stage after BTH application manifests itself through an increase in the expression level of DEGs and DEPs belonging to the WRKY family and GST family with high activity of protein kinases. In the presented studies, we also indicate the BTH impact on the activation of genes expression associated with the PTGS process in tomato plants. In the BTH-treated tomato plants, the regulation of the cellular detoxification process, which protects cells from apoptosis and DNA damage caused by ROS was activated. Moreover, for the first time, the BTH application impact on the thiamine metabolism pathway was presented in tomato plants, which may suggest that this vitamin plays a role in plants’ growth-defence balance in a tested experimental pathosystem. Also, changes in the amount level of I3CA and JA-Ile, together with up-regulation of the expression level of these hormones marker genes (*Solyc02g064830, Solyc12g009220*, and *Solyc08g036660*) may suggest that BTH is not only responsible for SAR induction but also activates other pathways, thus increasing the effectiveness of the plant defence against a broad spectrum of pathogens.

## Abbreviations

ABA: Abscisic acid
AUX/IAA: Auxin
BTH: Benzo(1,2,3)-thiadiazole-7-carbothioic acid S-methyl ester
DEG: Differentially expressed gene
DEP: Differentially expressed protein
GLM: Quasi Likelihood-F test
GO: Gene Ontology
KEGG: Kyoto Encyclopaedia of Genes and Genomes
HR: Hypersensitive response
I3CA: Indole-3-carboxilic acid
IAA: Indole-3-acetic acid
ISR: Induced systemic resistance
JA: Jasmonic acid
JA-Ile: Jasmonic-isoleucine
OPDA: Cis-(+)-12-oxo-phytodienoic acid
PR: Pathogenesis-related proteins
PTGS: Post-transcriptional gene silencing
RNA-Seq: RNA sequencing
SA: Salicylic acid
SAR: Systemic acquired resistance
TMM: Trimmed Mean of M values
ToMV: Tomato mosaic virus

## 5. Acknowledgments

The NGS RNA-Seq and targeted hormonal analysis were supported by the National Science Centre (Poland), project OPUS (No. UMO-2015/17/B/NZ9/01676) - “Systemic Acquired Resistance (SAR) of plants against viruses: new elicitors and biological and molecular characterization of their mechanism of action”. The Proteomic analysis has been supported by EPIC-XS, project number 823839, funded by the Horizon 2020 program of the European Union. The remaining analyses have been supported by the Ministry of Education and Science in Poland (No. Biotech-01).

## 6. Author contributions

A.OS., and P.F – conceptualisation. P.F. and A.OS. conceived and designed the study and wrote the manuscript with contributions from all coauthors. P.F. and P.W. performed the experiments. P.F., B.W., and A.OS. performed or supervised bioinformatic analysis of the RNA-Seq data. P.F., A.D., L.K., and A.OS. performed the protein isolation, identification, and bioinformatic analysis. P.F. and P.SB. performed the phytohormones analysis. A.OS. and P.F. analysed and interpreted all data. P.F., P.W., B.W., and A.OS. provided critical comments and edited the manuscript. All authors accepted the current form of the manuscript.

## 7. Conflict of interests

The authors declare no conflict of interest.

## 8. Data availability

All raw data generated in this study have been uploaded to the NCBI BioProject database under accession number PRJNA735633. The raw sequencing data of the transcriptome of each experimental variant were uploaded to the NCBI BioSample database under the accession numbers from SAMN19591107 to SAMN19591142. The mass spectrometry proteomics data were handled using the local laboratory information management system (LIMS) (Türker et al. 2010) and all relevant data have been deposited to the ProteomeXchange Consortium via the PRIDE (http://www.ebi.ac.uk/pride) partner repository with the data set identifier PXD028672.

## 10. Supplementary data

**Supplementary Table 1. (.pdf)** The summary results from the transcriptomic and proteomic analysis. A) The summary information of RNA-Seq analysis includes # of total reads and # of mapped reads received after the mapping process for each experimental condition; B) The summary information of # of total transcripts and proteins obtained after the analysis with # of up- and down-regulated DEGs and DEPs.

**Supplementary Table 2. (.xlsx)** All DEGs and DEPs (up- or down-regulated) obtained after analysis for each experimental condition.

**Supplementary Table 3. (.xlsx)** The 15 most enriched biological processes received for each experimental condition for transcriptomic and proteomic analysis separately.

**Supplementary Table 4. (.xlsx)** The 20 most regulated KEGG pathways received for each experimental condition for transcriptomic and proteomic analysis separately.

**Supplementary File 1. (.pdf)** Materials and Methods included A. Protein analysis protocol; B. Targeted hormonal analysis protocol and C. Table presenting information about the primers used in this publication.

## Notes

### Competing Interest Statement

The authors have declared no competing interest.

