## Supplemental File 1 for "Deciphering of BTH-induced response of tomato (*Solanum lycopersicum* L.) and its effect on plant virus infection through the multi-omics approach"

**Supplementary File 1**

**Materials and Methods**

### a. Protein analysis protocol

Fifty µg of protein were taken and reduced with 2 mM TCEP (tris(2-carboxyethyl)phosphine) and alkylated with 15 mM iodoacetamide at 60°C for 30 min. The protein concentration was estimated using the Qubit® Protein Assay Kit (Life Technologies, Switzerland).

The sp3 protein purification, digestion, and peptide clean-up were performed using a KingFisher Flex System (Thermo Scientific, USA) and Carboxylate-Modified Magnetic Particles (GE Life Sciences, USA)(Hughes *et al.*, 2014, Leutert *et al.*, 2019). Beads were conditioned following the manufacturer's instructions, consisting of 3 washes with water. Samples were diluted with an equal volume of 100% ethanol (50% ethanol final concentration). The beads, wash solutions, and samples were loaded into 96 deep well- or micro-plates and transferred to the KingFisher.

Following steps were carried out on the robot: a collection of beads from the last wash, protein binding to beads (14 min), washing of beads in wash solutions 1-3 (80% ethanol, 3 min each), protein digestion (for 4 h at 37°C with trypsin: protein ratio of 1:50 in 50 mM TEAB) and peptide elution from the magnetic beads (water, 6 min). The digest solution and water elution were combined and dried to completeness. Afterwards, the peptides were re-solubilized with 50 µl of 3% acetonitrile, 0.1% formic acid, and 3 µl of indexed retention time (iRT)-peptides (Biognosys, Switzerland) were spiked in each sample for MS analysis.

Mass spectrometry analysis was performed on an Orbitrap Fusion (Thermo Scientific, USA) equipped with a Digital PicoView source (New Objective, USA) and coupled to a M-Class UPLC (Waters, USA). The solvent composition of the two channels was 0.1% formic acid for channel A and 0.1% formic acid, 99.9% acetonitrile for channel B. For each sample, 3 µl of peptides were loaded on a commercial MZ Symmetry C18 Trap Column (100Å, 5 µm, 180 µm x 20 mm, Waters, USA) followed by nanoEase MZ C18 HSS T3 Column (100Å, 1.8 µm, 75 µm x 250 mm, Waters). The peptides were eluted at a flow rate of 300 nl/min. After an initial hold at 5% B for 3 min, a gradient from 5 to 24% B in 80 min and 36% B in 10 min was applied. The column was washed with 95% B for 10 min and afterwards the column was re-equilibrated to starting conditions for an additional 10 min. Samples were acquired in a randomized order. The mass spectrometer was operated in data-dependent mode (DDA) acquiring full-scan MS spectra (300−1,500 m/z) at a resolution of 120,000 at 200 m/z after accumulation to a target value of 400,000. Data-dependent MS/MS were recorded in the linear ion trap using quadrupole isolation with a window of 1.6 Da and HCD fragmentation with 35% fragmentation energy. The ion trap was operated in rapid scan mode with a target value of 8,000 and a maximum injection time of 80 ms. Only precursors with intensity above 5,000 were selected for MS/MS and the maximum cycle time was set to 3 s. Charge state screening was enabled. Singly, unassigned, and charge states higher than seven were rejected. Precursor masses previously selected for MS/MS measurement were excluded from further selection for 25 s, and the exclusion window was set at 10 ppm. The samples were acquired using internal lock mass calibration on m/z 371.1012 and 445.1200.

The acquired raw MS data were processed by MaxQuant (version 1.6.2.3), followed by protein identification using the integrated Andromeda search engine (Cox and Mann 2008). Spectra were searched against a tomato reference proteome (version from 2020-02-17), concatenated to its decoyed fasta database, and common protein contaminants. Carbamidomethylation of cysteine was set as fixed modification, while methionine oxidation and N-terminal protein acetylation were set as a variable. Enzyme specificity was set to trypsin/P allowing a minimal peptide length of 7 amino acids and a maximum of two missed cleavages. MaxQuant Iontrap default search settings were used. The maximum false discovery rate (FDR) was set to 0.01 for peptides and 0.05 for proteins. Label-free quantification was enabled and a 2 minutes window for the match between runs was applied. In the MaxQuant experimental design template, each file is kept separate in the experimental design to obtain individual quantitative values. Protein fold changes were computed based on Intensity values reported in the proteinGroups.txt file. A set of functions implemented in the R package SRMService (https://github.com/protViz/SRMService) was used to filter for proteins with 2 or more peptides allowing for a maximum of 4 missing values, and to normalize the data with a modified robust z-score transformation and to compute *p-Value* using the t-test with pooled variance. If all measurements of a protein are missing in one of the conditions, a pseudo fold change was computed replacing the missing group average by the mean of 10% smallest protein intensities in that condition.

In addition, we performed gene set enrichment analysis using the R/Bioconductor package fgsea (Korotkevich et al. 2016) and used gene sets specified in the molecular signature database (http://www.gsea-msigdb.org/). To apply GSEA to proteomics data, we mapped the UniProt identifiers to Entrez IDs using the UniProt mapping service. We ordered the protein lists using the log2FC. For cases where several UniProt IDs were mapped to a single Entrez ID, we averaged the log2FC.

**b. Targeted hormonal analysis protocol**

In brief, 30 mg of freeze-dried material of each sample condition were powdered and a pool of internal standards (abscisic acid-d6 (ABA-d6), salicylic acid-d5 (SA-d5), indole acetic acid-d4 (IAA-d4), dehydrojasmonic acid (dhJA), and jasmonic isoleucine-d6 (JA-Ile-d6) was added. Then hormone extraction was performed in a mixer mill. Samples were centrifuged and supernatant recovered and placed in a new tube to adjust the pH with acetic acid to 2.5-2.7. Two phases were created by adding diethyl ether and the organic phase was recovered, this step was repeated three times and the recovered organic phases were concentrated until dryness in a centrifugal evaporator (Speedvac). The extracted hormones were dissolved in 1 ml of MeOH/$H_2O$ with 0.01% HCOOH (10:90), resulting in a final concentration of internal standards of 100 ng/ml. External calibration curves of each pure standard were prepared for precise quantification.

The targeted hormonal analysis was performed in an Acquity ultraperformance liquid chromatography system (UPLC; Waters, Mildford, MA, USA) coupled to a triple quadrupole mass spectrometer (TQD, Waters, Manchester, UK). The column used for the LC separation was a UPLC Kinetex 2.6 µm EVO C18 100 Å, 2.1 x 50 mm (Phenomenex, USA). Conditions and solvent gradients used in this chromatographic analysis were the same as described in Sánchez-Bel et al. (2018) (Sánchez-Bel et al. 2018).

**c. Table presenting information about the primers used in this publication**

| Sequence ID | Primer Name | Primer seq. (5' - 3') | Description | Ta [˚C] | Length [nt] |
|---|---|---|---|---|---|
| *Solyc02g080890.3.1* | **Forward** | *GAAAGCTCGTGTCTCCGTTC* | ***WRKY transcription factor 6*** | 58 | 229 |
| | **Reverse** | *TGGTAATGGGTGGTTGTGTG* | | | |
| *Solyc08g062490.3.1* | **Forward** | *AGTGCAAACATGTTCGATGG* | ***WRKY transcription factor 50*** | 58 | 237 |
| | **Reverse** | *GCCAACGTTGTTGCTAGGTT* | | | |
| *Solyc04g081630.2.1* | **Forward** | *CAGGGTTGATTTCGAGCACT* | ***Heat shock protein 90 (AHRD V3.3 *-* Q9MB32_ORYSA)*** | 55 | 218 |
| | **Reverse** | *CAAATTCCACGAGGAAGACC* | | | |
| *Solyc07g064160.3.1* | **Forward** | *TGTCATTGTTGGTGCTGGAT* | ***Thiamine thiazole synthase, chloroplastic (AHRD V3.3 *** K4CH99_SOLLC)*** | 58 | 150 |
| | **Reverse** | *CGCACAACCATAGCAGAGAA* | | | |
| *Solyc09g011500.3.1* | **Forward** | *TGAAGGCCCTTACATCTTGC* | ***Glutathione S-transferase-like protein (AHRD V3.3 *** K7VK72_SOLTU)*** | 55 | 214 |
| | **Reverse** | *TCCAAATTTGTCACCCACAA* | | | |
| *Solyc07g056510.3.1* | **Forward** | *TCCCTTTGGTATGAGGGTGA* | ***Glutathione S-transferase (AHRD V1 **** D3Y4H6_9ROSI)*** | 58 | 213 |
| | **Reverse** | *AGTGGAGCTTTGTCCTTCCA* | | | |
| *Solyc10g018340.1.1* | **Forward** | *AAGTGTGGTGCCAAAAGGTC* | ***Small auxin up-regulated RNA71*** | 58 | 182 |
| | **Reverse** | *CCGAGATCCGACAAGGAATA* | | | |
| *Solyc02g064830.3.1* | **Forward** | *CAGTGCCGAGTTGCAGATAA* | ***Indole-3-acetic acid-amido synthetas 3-3*** | 58 | 233 |
| | **Reverse** | *TGTAAACGCTGTCGTTCTGG* | | | |
| *Solyc12g009220.2.1* | **Forward** | *CGTCCGTTGAAACAAATCCT* | ***Jasmonate ZIM-domain protein 1*** | 58 | 218 |
| | **Reverse** | *GGGGTTCTGTTTGTTGGCTA* | | | |
| *Solyc08g036660.3.1* | **Forward** | *GAGCAGAAATCTGAGCCGTTA* | ***Jasmonate zim-domain protein 11 (AHRD V3.3 *** G7IP70_MEDTR)*** | 58 | 240 |
| | **Reverse** | *TGGCGAAGTTGTTTGAGTTCT* | | | |
| *Solyc06g071050.3.1* | **Forward** | *CTGGCAGACAAAGCAAATGA* | ***Hypersensitive-induced response protein (AHRD V3.3 *** A6YGE4_CARPA)*** | 50 | 180 |
| | **Reverse** | *AGCCGACATAGCCTTCTCAA* | | | |
| *Solyc02g092580.3.1* | **Forward** | *GACCCAACCCTTAACAAGCA* | ***Peroxidase (AHRD V3.3 *** K4BD54_SOLLC)*** | 50 | 218 |
| | **Reverse** | *TTACTGGCCCAAAGATCCAC* | | | |
| *Solyc01g008620.3.1* | **Forward** | *GGTCTCAACCGCGACATATT* | ***Beta-1,3-glucanase (AHRD V3.3 *** Q9SYX6_TOBAC)*** | 56 | 170 |
| | **Reverse** | *TATCATCAGCATGGCCAAAA* | | | |
| *Solyc02g069260.3.1* | **Forward** | *CCGACCATGACTTTCGTTTT* | ***Argonaute2a*** | 56 | 199 |
| | **Reverse** | *TTGCAGCTCGAATGTTGTTC* | | | |
| *Solyc03g123800.2.1* | **Forward** | *CGTGAGGTCAAGATTGCAGA* | ***MAP kinase kinase 2*** | 56 | 158 |
| | **Reverse** | *CATATGTCCCCAGCATACCC* | | | |
| *Solyc07g043480.1.1* | **Forward** | *AGTCGCGGAAGTGCTAAAAA* | ***Glycosyltransferase (AHRD V3.3 *** A0A0A1WC49_NICAT)*** | 56 | 166 |
| | **Reverse** | *CGCTTTACAGCCGTCCTTAG* | | | |
| *Solyc03g111610.3.1* | **Forward** | *TTCAATGGCGTTGAGAGTTG* | ***Haloacid dehalogenase-like hydrolase (HAD) superfamily protein (AHRD V3.3 *** AT3G48420.1)*** | 56 | 193 |
| | **Reverse** | *GCCTACTCGTCTTGCTACGG* | | | |
| *Solyc04g076790.3.1* | **Forward** | *GCGCTAGATCTTCCTCATGG* | ***Serine hydroxymethyltransferase (AHRD V3.3 *** M1A7K0_SOLTU)*** | 56 | 202 |
| | **Reverse** | *GAGCATAAGCACTTGCACCA* | | | |
| *Solyc06g005060.3.1* | **Forward** | *ATTGGAAACGGATATGCTCCA* | ***Elongation factor 1-alpha*** | 55 | 110 |
| | **Reverse** | *TCCTTACCTGAACGCCTGTCA* | | | |
| *Solyc11g005330.1.1* | **Forward** | *TGTCCCTATTTACGAGGGTTATGC* | ***Actin (AHRD V1 ***- Q7XZK0_GOSHI)*** | 55 | 76 |
| | **Reverse** | *CAGTTAAATCACGACCAGCAAGAT* | | | |
