## Supplemental Table 1 for "Deciphering of BTH-induced response of tomato (*Solanum lycopersicum* L.) and its effect on plant virus infection through the multi-omics approach"

**Supplementary Table 1. The summary results from the transcriptomic and proteomic analysis. a) The summary information of RNA-Seq analysis includes # of total reads and # of mapped reads received after the mapping process for each experimental condition; b) The summary information of # of total transcripts and proteins obtained after the analysis with # of up- and down-regulated DEGs and DEPs.**

**a)**

| *Experimental Variants* | *Total reads (4n)* | *Mapped Reads* | | *Aligned Reads* | | | |
|---|---|---|---|---|---|---|---|
| | | *Uniquely Mapped Reads (4n)* | *Reads Mapped to Multiple Loci (4n)* | *Feature* | *No Feature* | *Ambiguous* | *Alignment Not Unique* |
| *Control 1dpt* | 62,146,164 | 59,712,632 / 96.10% | 966,054 / 1.55% | 48,012,027 / 77.26% | 2,297,369 / 3.70% | 9,403,236 / 15.13% | 2,362,330 / 3.62% |
| *BTH 1dpt* | 62,663,391 | 59,939,122 / 95.65% | 1,220,259 / 1.95% | 47,476,058 / 75.76% | 2,348,785 / 3.75% | 10,114,279 / 16.14% | 3,019,209 / 4.82% |
| *Control 8dpt* | 66,743,475 | 64,773,333 / 97.05% | 737,726 / 1.11% | 53,269,948 / 79.81% | 2,563,828 / 3.84% | 8,939,557 / 13.39% | 1,773,656 / 2.66% |
| *BTH 8dpt* | 63,795,529 | 61,688,343 / 96.70% | 770,607 / 1.21% | 50,785,017 / 79.61% | 2,625,628 / 4.12% | 8,277,698 / 12.98% | 1,876,378 / 2.94% |
| *BTH-ToMV* | 64,058,933 | 62,336,515 / 97.31% | 634,276 / 0.99% | 52,015,160 / 81.20% | 2,546,625 / 3.98% | 7,774,727 / 12.14% | 1,530,011 / 2.39% |
| *ToMV* | 66,136,747 | 64,079,633 / 96.90% | 728,728 / 1.10% | 53,198,019 / 79.81% | 2,312,883 / 3.50% | 8,568,731 / 12.96% | 1,728,147 / 2.61% |

**b)**

| *Experimental Variants* | *Annotated transcripts* | *DEGs (p.Val=0.05, -1 ≥ logFC ≥ 1)* | | *SUM of DEGs* | *Annotated Proteins* | *DEPs (p.Val=0.05, -0.75 ≥ logFC ≥ 0.75)* | | *SUM of DEPs* |
|---|---|---|---|---|---|---|---|---|
| | | *UP-regulated* | *Down-regulated* | | | *UP-regulated* | *Down-regulated* | |
| *BTH 1dpt* | 21,834 | 1469 | 1173 | 2642 | 3459 | 124 | 59 | 183 |
| *BTH 8dpt* | | 899 | 455 | 1354 | 3562 | 287 | 228 | 515 |
| *BTH-ToMV* | | 984 | 1127 | 2111 | 3559 | 195 | 121 | 317 |
| *ToMV* | | 140 | 170 | 310 | 3460 | 51 | 105 | 156 |
